# Invasive genetic rescue: Dispersal following repeated culling reinforces the genetic diversity of an invasive mammal

**DOI:** 10.1101/2022.06.09.495496

**Authors:** Rebecca Synnott, Craig Shuttleworth, David J. Everest, Claire Stevenson-Holt, Catherine O’Reilly, Allan D. McDevitt, Denise B O’Meara

**Affiliations:** Molecular Ecology Research Group, Eco-Innovation Research Centre, Department of Science, South East Technological University (SETU), Cork Road, Waterford, Ireland; School of Natural Sciences, Bangor University, Bangor, Gwynedd, Wales, United Kingdom (UK); Animal and Plant Health Agency (APHA)-Weybridge, Department for Environment, Food and Rural Affairs, London, UK; Department of Science, Natural Resources and Outdoor Studies, University of Cumbria, Ambleside, UK; Department of Natural Sciences, School of Science and Computing, Atlantic Technological University, Galway, Ireland

## Abstract

Since its introduction from the United States in 1876 the invasive North American Eastern grey squirrel (*Sciurus carolinensis*) has contributed to the decline of the native Eurasian red squirrel (*Sciurus vulgaris*) in Britain. Consequently, grey squirrel populations are often subjected to extensive control efforts in order to reduce local abundance and prevent spread, particularly within habitats containing red squirrels. Grey squirrel removal from the island of Anglesey off the coast of north Wales began in 1998 and was completed in 2013. A parallel successful red squirrel reinforcement translocation project also took place. The narrow sea-channel, road and rail bridge connection has meant that the adjacent mainland grey squirrel population has been controlled in subsequent years to reduce the probability of re-invasion. The aim of this study was to assess the overall impact of repeated culling efforts carried out between 2011 and 2020 on the genetic diversity of the grey squirrel population in north Wales to inform future adaptive management plans. Using a combination of mitochondrial DNA (mtDNA) and microsatellite DNA analysis, we found high genetic diversity in both marker types, with six diverse mtDNA haplotypes found and relatively high levels of nuclear genetic diversity, even after repeated culling efforts. Our results suggest that ongoing culling efforts may not adequately reduce genetic diversity to a level where it contributes to a long-term population decline.

## Introduction

Global human impacts on the natural environment are jeopardising species and ecosystems at ever increasing rates (Tilman et al 2017). These impacts occur from land use changes, the fragmentation of habitats, overhunting and exploitation of wildlife populations and the introduction and spread of invasive species (Tilman et al 2017). Invasive species represent one of the greatest threats to loss of biodiversity (Bellard et al 2016). The success of a particular invasion depends on local habitat diversity and ecosystem resilience which ultimately determines whether a species will become invasive (Williamson 2006; Hayes & Barry 2008). Successful invasions can have negative ecological and economic affects, and it is these species that cause conservation concern requiring resources to control their spread (Manchester & Bullock 2000). Eradication programmes and ongoing biosecurity surveillance of invasive species are costly for both government and non-government organisations (Olson 2006). However, it is noteworthy that prolonged invasive species control efforts can lead to a loss of genetic diversity amongst residual invasive populations, an effect that can accelerate population decline and ultimately increase the success of an eradication programme (Roman & Darling, 2007; Zalewski et al 2016; Browett et al 2020).

The presence of the North American Eastern grey squirrel (*Sciurus carolinensis*) in Britain is a significant threat to forest ecosystems (Lowe et al 2010; Shuttleworth et al 2016). Grey squirrels were introduced into Britain in the late 19^th^ and early 20^th^ centuries, with at least eight documented introductions including a minimum of 135 recorded individuals from the USA and Canada (Shorten 1954). These founding individuals introduced for ornamental purposes, quickly established, and subsequently spread to further locations or were chosen for some of the additional 31 documented translocations that took place across the country between 1876 and 1937 (Middleton 1931; Shorten 1954). Despite ecological and climatic differences between North America and Britain, the founding individuals often led to the establishment of large, viable populations (Gurnell et al 2004; Stevenson-Holt & Sinclair 2015). Mitochondrial DNA (mtDNA) studies show that there is a high diversity of haplotypes found in Britain today, which reflects the large number of original introductions from a broad geographic area, confirming that the contemporary population has been derived from multiple sources (David-Gray et al 1998; Stevenson et al 2013; Stevenson-Holt & Sinclair 2015). Nuclear or microsatellite DNA has shown that high levels of genetic differentiation coupled with low levels of migration and regional gene flow occur within the British grey squirrel populations (Signorile et al 2016).

At the time of the grey squirrel introduction, the native Eurasian red squirrel (*S. vulgaris*) population had been in a state of decline since the 1700s due to harsh winters, deforestation associated with industrialization and shipbuilding, hunting and the trade of wild animals, with red squirrels particularly popular. By the late 19^th^ Century, there were over 20,000 red squirrels sold annually in London markets, and such was the demand that red squirrels were imported from throughout Europe due to local shortages (Shorten 1954; O’Meara et al 2018). Grey squirrels contributed to further red squirrel population declines as a result of resource competition and the spread of squirrelpox, which is asymptomatic in grey squirrels but pathogenic in red squirrels (Chantrey et al 2014; McInnes et al 2020). As a result of regional population extinction in the face of grey squirrel spread, the red squirrel is of conservation concern within Britain. Conservation efforts to restore and enhance populations are widespread (Shuttleworth, Lurz & Robinson 2021) including the island of Anglesey off the coast of north Wales, which is connected to the mainland via two man-made bridges (Ogden et al 2005).

Complete eradication of grey squirrels from Anglesey occurred in the summer of 2013 (Schuchert et al 2014; Shuttleworth et al 2015), but two years later, several grey squirrels were reported by local people, requiring reactive management by the local conservation team. Nuclear DNA was used to identify genetic similarities between the Anglesey grey squirrel population and those on the adjacent mainland indicating that animals had naturally colonised Anglesey, although some genetically differentiated animals were also found in the southwest of the island, suggesting a separate colonisation event had also occurred. Reinvasion was likely to have occurred from the mainland by grey squirrels swimming the sea-channel, crossing one of the bridges, or inadvertently/intentionally transported in vehicles and subsequently released (Signorile & Shuttleworth 2016; Shuttleworth 2021). Consequently, the long-term maintenance of red squirrels on Anglesey is reliant on the continued absence of grey squirrels from the island, requiring continuous mainland control efforts (Shuttleworth & Halliwell (2016); Shuttleworth et al 2020).

The aim of this study was to genetically assess culled grey squirrels in north Wales over a period of nine years, with a view to understanding how the eradication efforts have impacted the contemporary diversity of the species over time. We also aimed to investigate the recolonization of the species following eradication, while also considering the genetic ancestry and legacy of the species’ original introductions from North America. Based on our findings, we make suggestions regarding the long-term management of the grey squirrel population in north Wales with a view to maintaining the red squirrel population on Anglesey.

## Materials and Methods

### Ethical statement

Grey squirrels were culled by trained practitioners as part of ongoing efforts to manage and reduce the invasive population. Animals were trapped and humanely dispatched as per Schedule 9 of the Wildlife and Countryside Act (Wildlife and Countryside Act 1981, 2017), which makes it illegal to release or allow invasive species to escape into the wild. UK Best Practice was adopted and included high welfare standards (Gill et al 2019). Animals were not specifically killed for this study, but samples were opportunistically retained over the course of trapping efforts in the event that potential studies may arise where such samples might be of interest.

### Sample collection

A total of 410 samples, 341 tissue and 69 hair samples were collected between 2011 and 2020 by the Red Squirrel Trust Wales (RSTW). Grey squirrel trapping was conducted in woodland areas on Anglesey (ANG), in Gwynedd (GWY) and Clocaenog (CL) Forest (Fig. 1). A tissue or hair sample (30-40 tail hairs with follicles) was taken from each culled individual and stored at -20 °C.

**Figure 1.**
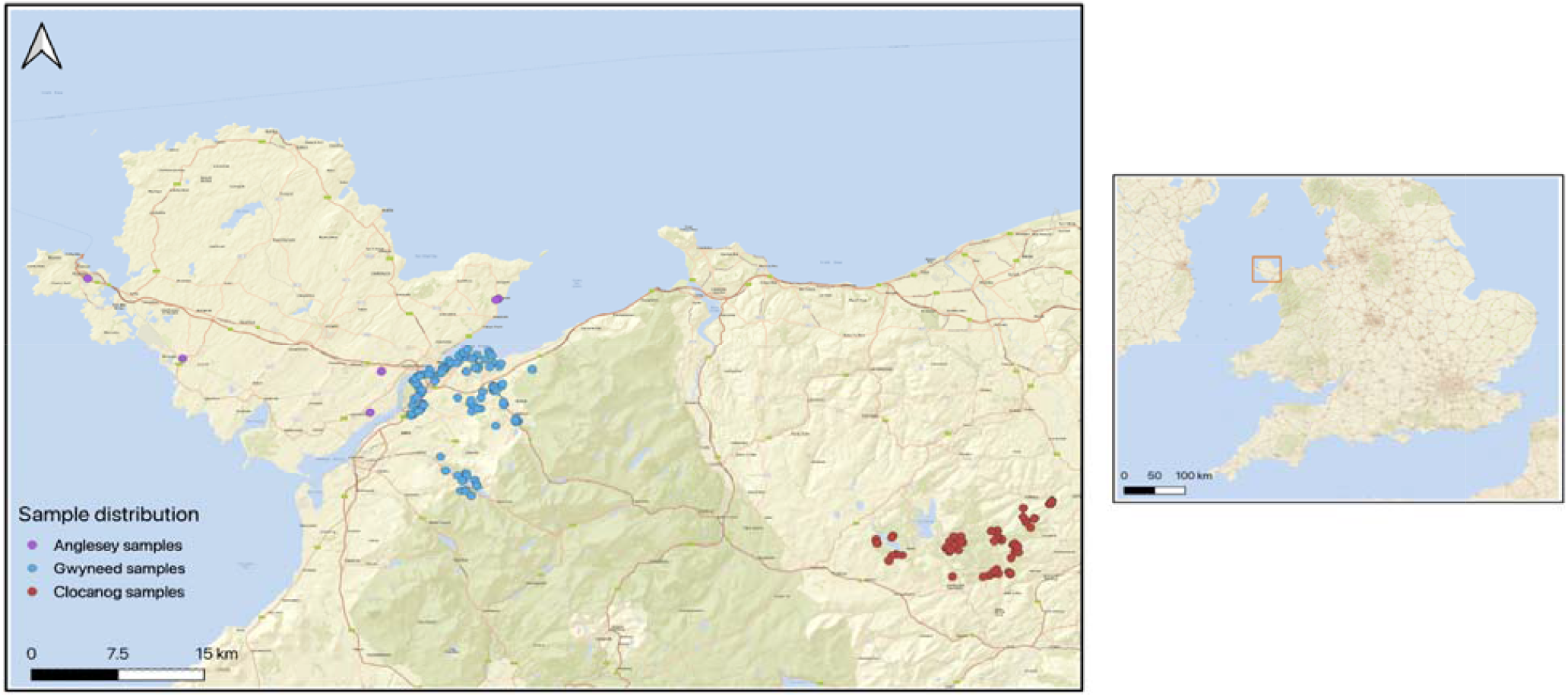
Left inset: The distribution of grey squirrel samples collected from three areas (Anglesey, Gwynedd and Clocaenog) across north Wales. Right inset: the location of north Wales, highlighted by the red square, relative to southern Britain, the east of Ireland and the north of France.

### DNA Extraction

Genomic DNA (gDNA) was extracted from tissue samples using the QIAGEN DNeasy Blood and Tissue® kits as per the manufacturer’s instructions. The hair samples were extracted using the Macherey-Nagel™ NucleoSpin™ Tissue kit following the instructions provided for hair samples. The quality and quantity of the extracted tissue DNA was determined using the Nanodrop™ 8000 (Thermo Fisher Scientific Inc., USA). Purified DNA preparations were stored at -20 °C.

### Mitochondrial DNA analysis

MtDNA analysis was performed on all samples extracted as part of this study. D-loop mtDNA primers published by Stevenson-Holt et al (2013) were used to amplify a 329-bp region using the follow primer pair: Dloop forward primer 5’-GCCACCCCCAAGTTAAATGG-3’ and Dloop reverse primer 5’-ATTCGTGCATTAATGCACTATCC-3’. A 10 μl PCR reaction mix consisted of 5 μl GoTaq® Hot Start Green Master Mix (Promega), 1 μl of a primer mix containing 5 μm each of the forward and reverse primers, 3 μl of molecular grade water and 1 μl of the DNA extract. Thermocycling conditions consisted of denaturation step of 1 min at 94 °C followed by 40 cycles of 30 s at 94°C, annealing 30 s at 52 °C, extension 1 min at 72 °C, final extension of 5 min at 72 °C. PCR products were visualised using gel electrophoresis and positive PCR products were cleaned using the *micro*Clean (Clent Life Science, Stourbridge, UK) and sequenced using the forward primer on the 3500 Genetic Analyzer (Applied Biosystems).

### MtDNA data analysis

MtDNA sequences generated in this study were compared by multiple alignments using the CLUSTALW method in MEGA, version 7 (Tamura et al 2011). Haplotypes were identified using ARLEQUIN, version 3.5 (Excoffier and Lischer 2010). MtDNA haplotypes from this study were compared to those previously recorded in Britain (N = 12) by Stevenson-Holt & Sinclair (2015). An additional 50 mtDNA grey squirrel sequences (486 bp) sampled in North America and published by Moncrief et al (2008) (accession numbers JX104415–JX104465) were compared to the British samples by multiple alignments using the CLUSTALW method as previously outlined. Data included the haplotypes previously identified by Stevenson-Holt & Sinclair (2015) and were combined with the data from this study and grouped as follows: Eastern USA (Virginia and Maryland) (N = 20), Mid-Western USA (Indiana) (N = 3) and Southern USA (Alabama, Louisiana, Mississippi, Georgia and Tennessee) (N = 27). Using 328 bp of the d-loop, a TCS network was constructed using a parsimony approach (Clement et al 2002) in the programme POPART v.1.7.1 (Leigh & Byrant 2015), and haplotypes were colour coded by region of origin.

### Microsatellite DNA analysis

A subset of the samples that were successfully haplotyped were subsequently genotyped (N = 256) using the following microsatellite loci: SCV3, SCV4 SCV6, SCV9, SCV15 (Hale et al 2001), LIS3 (Shibata et al 2003) GR05, GR11 (Fike et al 2013), FO11 (Fike & Rhodes 2009) (Table 1). Samples were genotyped in duplicate using three multiplex reactions.

**Table 1.**
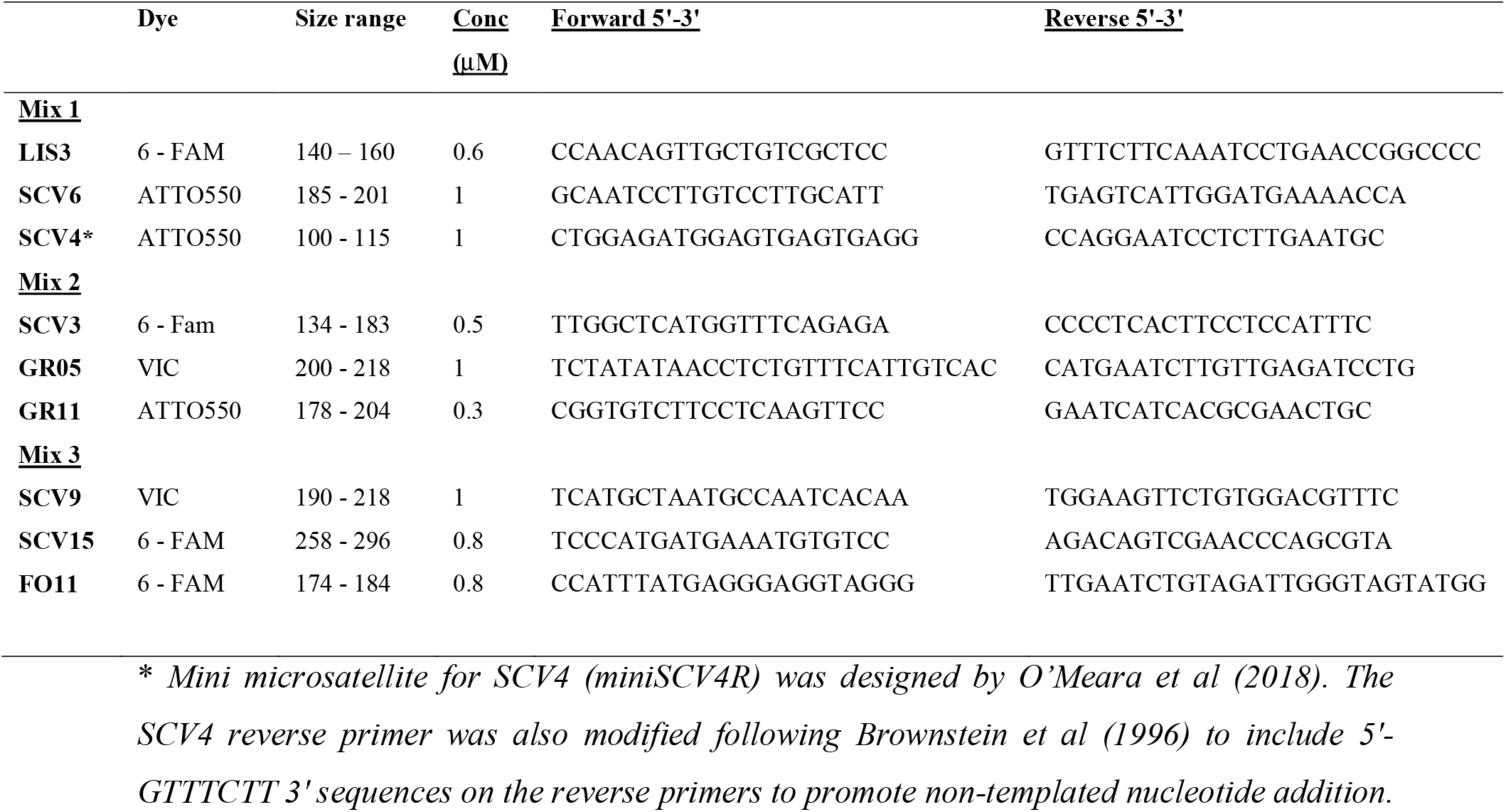
Details of nine microsatellite loci used in this study: SCV3, SCV4 SCV6, SCV9, SCV15 (Hale et al 2001), LIS3 (Shibata et al 2003) GR05, GR11 (Fike et al 2013), FO11 (Fike and Rhodes 2009). Other details provided include the three PCR multiplexes, the fluorescent dyes, expected size range and the concentration of each primer added to the multiplex reaction.

**Table 2:** Descriptive statistics for grey squirrels in north Wales across nine microsatellite loci.

|  | SCV4 | LIS3 | SCV6 | SCV3 | GR11 | GR05 | SCV15 | SCV9 | FO11 | Average |
| --- | --- | --- | --- | --- | --- | --- | --- | --- | --- | --- |
| **N** | 255 | 256 | 255 | 252 | 250 | 240 | 255 | 255 | 241 | 251 |
| **N<sub>a</sub>** | 7 | 6 | 7 | 1 | 8 | 4 | 11 | 7 | 9 | 6.67 |
| **A<sub>R</sub>** | 7 | 5.9 | 7 | 1 | 7.9 | 4 | 10.8 | 6.9 | 9 | 6.61 |
| **H<sub>O</sub>** | 0.706 | 0.102 | 0.443 | 0.000 | 0.400 | 0.171 | 0.549 | 0.431 | 0.382 | 0.35 |
| **H<sub>E</sub>** | 0.767 | 0.496 | 0.794 | 0.000 | 0.694 | 0.266 | 0.757 | 0.677 | 0.606 | 0.56 |
| **H<sub>eq</sub>** | 0.677 | 0.627 | 0.679 | 0.000 | 0.717 | 0.477 | 0.796 | 0.677 | 0.677 | 0.59 |
| **F<sub>IS</sub>** | **0.085** | **0.795** | **0.44** | 0 | **0.422** | **0.36** | **0.272** | **0.372** | **0.37** | 0.35 |

**Table 3:** Average descriptive statistics for the grey squirrel populations in north Wales across eight loci. Abbreviations are as follows: number of samples amplified per loci (N), number of alleles per loci (A), Allelic Richness (A_R_) observed heterozygosity (H_O_), expected heterozygosity (H_E_), expected heterozygosity at equilibrium (H_EQ_), inbreeding coefficient (F_IS_). Values in bold indicate significant deviation from Hardy-Weinberg Equilibrium at P = 0.05, and also after Bonferroni correction P = 0.00078.

| SITE | YEAR | N | A | $A_R$ | $H_O$ | $H_E$ | $H_{EQ}$ | $F_{IS}$ |
| --- | --- | --- | --- | --- | --- | --- | --- | --- |
| ANGLESEY | 2015 | 3 |  |  |  |  |  |  |
|  | 2016 | 1 |  |  |  |  |  |  |
|  | 2017 | 2 |  |  |  |  |  |  |
|  | 2020 | 1 |  |  |  |  |  |  |
| GWYNEDD | 2014 | 68 | 5.9 | 4.59 | 0.410 | 0.612 | 0.645 | **0.337** |
|  | 2016 | 26 | 5.1 | 4.43 | 0.409 | 0.589 | 0.646 | **0.325** |
|  | 2017 | 30 | 5 | 4.44 | 0.513 | 0.624 | 0.622 | **0.193** |
|  | 2019 | 28 | 4.9 | 4.37 | 0.353 | 0.609 | 0.610 | **0.436** |
| CLOCAENOG FOREST | 2011 | 15 | 4 | 3.90 | 0.336 | 0.494 | 0.573 | **0.350** |
|  | 2012 | 23 | 4 | 3.75 | 0.366 | 0.533 | 0.520 | **0.334** |
|  | 2014 | 18 | 4.3 | 4.01 | 0.382 | 0.595 | 0.580 | **0.382** |
|  | 2015 | 39 | 5 | 4.07 | 0.372 | 0.551 | 0.620 | **0.336** |

The PCR reaction contained 5 μl GoTaq® Hot Start Green Master Mix (Promega), 1 μl of the DNA, primer mix containing 5 μm each of the forward and reverse primers (concentration specified in Table 1) and molecular grade water in a 10 μl reaction. Thermocycling conditions consisted of an initial step of 95 °C for 10 min, followed by 20 cycles of 95 °C for 30 sec and a touchdown from 65 to 55 °C for 1 min decreasing by 0.5 °C per cycle, and then 72 °C for 1.5 min. This was followed by 20 cycles of 95 °C for 30 sec, 55 °C for 1 min, 72 °C for 1.5 min and a final extension of 72 °C for 10 min. Fragment analysis was completed using the 3500 Genetic Analyzer (Applied Biosystems). Alleles were scored using the GeneMapper software Version 5 (Applied Biosystems). All samples were amplified in duplicate and failed or inconsistent scores across both replicates were independently repeated from the PCR stage.

### Microsatellite data analysis

The dataset was assessed for the presence of errors, including scoring error due to stuttering, allele dropout and the presence of null alleles, using the program MICRO-CHECKER v.2.2.3 (van Oosterhout et al 2004). As samples consisted of a mixture of tissue and hair sourced DNA, the dataset was assessed to ensure it consisted of unique individuals and that the microsatellite panel used was sufficient to identify animals at an individual level. GENALEX v.6.5b (Peakall & Smouse (2006) was also used to assess probability of identify (PI) and probability of sibship (PIsib), i.e. to assess how many microsatellites were required to identify unique, unrelated individuals, and related individuals.

Descriptive statistics were calculated for the overall sample set, and also repeated by year and area of cull to establish the genetic consequences of repeated culling. GENALEX was used to calculate observed (H_O_) and expected (H_E_) heterozygosity, the number of alleles (N_A_). Allelic Richness (A_R_) and the inbreeding coefficient (F_IS_) was calculated using FSTAT version 2.9.3 (Goudet 1995) with significance levels for F_IS_ levels calculated by randomizing the alleles among the individuals within the population and comparison to the observed data to determine deviations from Hardy-Weinberg Equilibrium using 10,000 permutations. Tests for linkage disequilibrium were performed between pairs of loci using GENEPOP v.4.7 (Raymond & Rousset 1995; Rousset, 2008)

The program BOTTLENECK v.1.2.02 (Piry et al 1999) was used to assess the dataset for evidence of a genetic bottleneck. Bottlenecks are detected when there is an excess of heterozygosity in comparison to the number of alleles, as the number of alleles declines in a population before there is an impact on the level of heterozygosity (Luikart & Cornuet (1998). This test was carried out using the two-phase model (TPM), which is recommended for use with microsatellite data. The TPM was used with the following settings: 80% single-step mutations, a variance among multiple steps of 12, and 5,000 iterations. The probability of significant heterozygosity excess was subsequently determined using Wilcoxon’s signed rank test and the mode-shift indicator test which can detect the presence of a recent genetic bottleneck. A mode found outside of this normal L-shaped distribution is transient and detectable if the bottleneck has occurred in the last few dozen generations (Luikart et al 1998).

STRUCTURE 2.3.4. (Pritchard et al., 2000) was used to assign individuals to clusters using a Bayesian approach. A burn-in period of 250,000 Markov chain Monte Carlo steps was selected, followed by length of 750,000. An admixture model was chosen with LOCIPRIOR. The K values that were selected were 1 – 10 with the number of iterations set to 5. The results file generated was also entered into STRUCTURE Harvester (Earl & vonHoldt 2012) to assess and visualise all likelihood values across all values of K. The web server CLUMPAK was used to summarise and visualise the STRUCTURE results (Kopelman et al 2015). To further investigate the presence of genetic structure, a principal coordinate analysis (PCoA) was generated in GENALEX and visualised using ggplot in R.

## Results

### MtDNA

A total of six mtDNA haplotypes were recorded from 340 successfully sequenced individuals sampled across north Wales and Anglesey, three of which were new to this study, named H13, H14 and H15, following on from the 12 haplotypes described by Stevenson-Holt & Sinclair (2015) (Fig. 2). Three of the haplotypes H7, H9 and H12 were previously identified by Stevenson-Holt & Sinclair (2015). All six haplotypes were found in Clocaenog, five in Gwynedd (excluding H15) and three in Anglesey (including H9, H12 and H14). When the haplotypes recorded in north Wales were compared to the haplotypes previously recorded in Britain by Stevenson-Holt & Sinclair (2015), H7, H9 and H12 had been previously recorded in northern England including Cheshire, Lancashire and Cumbria.

**Figure 2.**
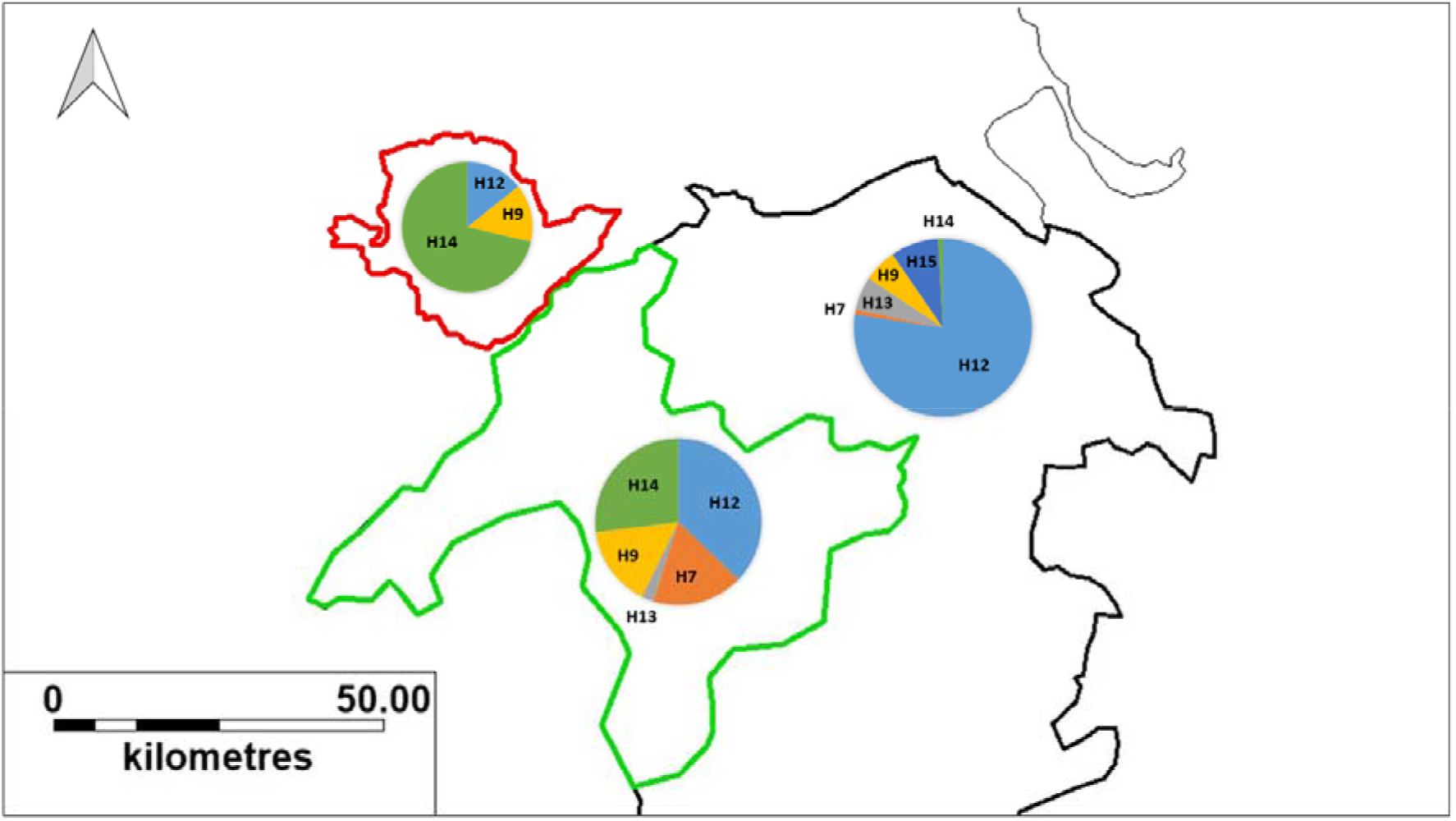
Distribution of mtDNA haplotypes found in Anglesey (N = 7) (location border in red), Gwynedd (N = 217) (border in green) and Clocaenog (N = 116) (border in black).

The haplotypes found within Britain were also compared to those recorded in the USA by Moncrief et al (2012) (Fig. 3). The Network diagram was split into two clusters, with some of the British haplotypes found in both clusters. The haplotypes from this study were from the same cluster, but were distributed throughout. H15 and H9 were clustered on the same branch, indicating some genetic similarity, and some of the British haplotypes (H8, H11 and H12) were also recorded in the Eastern US states.

**Figure 3.**
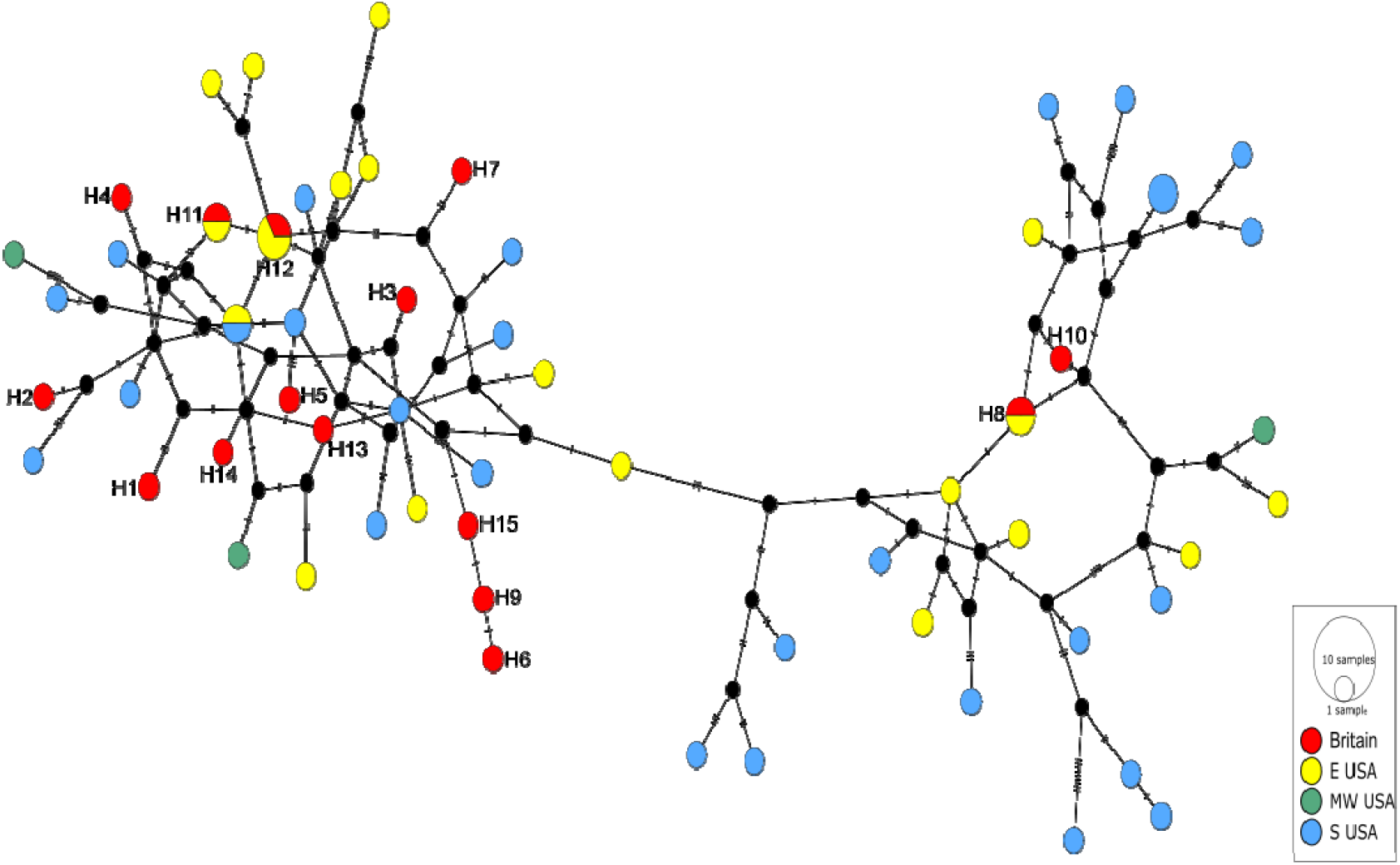
TCS network diagram generated using 328 bp D-loop mitochondrial DNA combing the new sequences generated from this study (H13, H14 and H15) with those previously published from Stevenson-holt et al (2015) and Moncrief et al (2012). The British haplotypes H1 – H15 have been labelled for clarity.

### Microsatellite

Of the 256 successfully genotyped individuals, the success averaged 98% across all loci, ranging from 95.6% at GR05 to 100% at LIS3. No evidence of large allele dropout was detected at any loci, but evidence of homozygote excess was detected at the following loci: SCV9, SCV15, GR11, SCV6 and LIS3 (P < 0.05). Although an excess of homozygotes can be caused by stuttering or the presence of null alleles, we carefully re-examined all data which had been analysed in duplicate from the PCR stage for consistency and found no evidence of human error. Four animals were found to have two matching genotypes; C15166 and G1653 were found to be identical as were C1217 and C15137. In both cases animals were successfully genotyped across all loci. Probability of identity averaged 2.9 × 10^−1^ across all loci, with a cumulative PI of 4.5 × 10^−7^. Probability of sibship averaged 5.4 × 10^−1^ across all loci, with a cumulative PIsib of 2.4 × 10^−3^. The number of alleles average 6.7, with similar levels of allelic richness observed at 6.6. Observed heterozygosity averaged 0.35, while expected heterozygosity averaged 0.56. F_IS_ values averaged 0.35 and all loci deviated significantly from Hardy–Weinberg equilibrium at the 5% significance level. SCV3 showed no variability across the dataset. Linkage disequilibrium was detected in eight of the 36 possible pairwise comparisons at the 5% significance level, three of which remained significant following a Bonferroni correction (P = 0.0014). This included the following pairs: SCV6 and GR05; SCV4 and SCV15 and SCV4 and FO11. No consistent patterns were observed (Tab. 2).

There was no evidence of a genetic bottleneck in the dataset. The data showed that while some loci exhibited differences in H_E_ and H_EQ_, there was not a significant excess of heterozygosity relative to the number of alleles due to differences between the levels of expected heterozygosity (e) relative to heterozygosity estimated under mutation–drift equilibrium (H_EQ_) (Tab. 2). The probability value for a one-tailed Wilcoxon’s signed rank test for an excess of heterozygosity was P = 0.84. The mode-shift test also showed a normal L-shaped distribution providing further evidence for a lack of a genetic bottleneck.

The calculations for genetic diversity and genetic bottlenecks were repeated by dividing the data by locality and year of cull (Tab. 3). The number of animals culled in Anglesey ranged from one in 2016 and 2020 to three animals in 2014. Due to the small population sizes, further calculations were not completed for this site. Culling took place in Gwynedd and Clocaenog between 2011 and 2019 with sampling sizes ranging between 15 and 68. Overall, the number of alleles and allelic richness remained similar across all years and localities, although a slight decrease was observed in Gwynedd (A_R_ = 4.59 - 4.37). Levels of expected and observed heterozygosity also remained relatively consistent in Clocaenog, with a slight increase in both statistics occurring throughout the cull period (H_O_ = 0.494 -0.551; H_E_ = 0.494 – 0.551), while in Gwynedd, a slight overall decrease occurred (H_O_ = 0.410 - 0.353; H_E_ = 0.612 – 0.609). F_IS_ values were positive and significant at the 5% level for all populations indicated high levels of inbreeding. No significant evidence of population bottlenecks occurred.

### Genetic structure

A total of 254 individuals were analysed for population based assignment. The delta K or Evanno method showed that K = 2 was the most likely number of genetic clusters within the population (Fig. 4A) (Evanno et al 2005). The plot of the mean likelihood, L(K), established from combining each replicate per K value and associated standard deviation from STRUCTURE HARVESTER showed a gradual plateau occurring in the dataset from K = 3 to K = 5, but this also coincided with an increase in variance suggesting the true K value lay between K = 2 or K = 5 (Fig. 4B).

**Figure 4:**
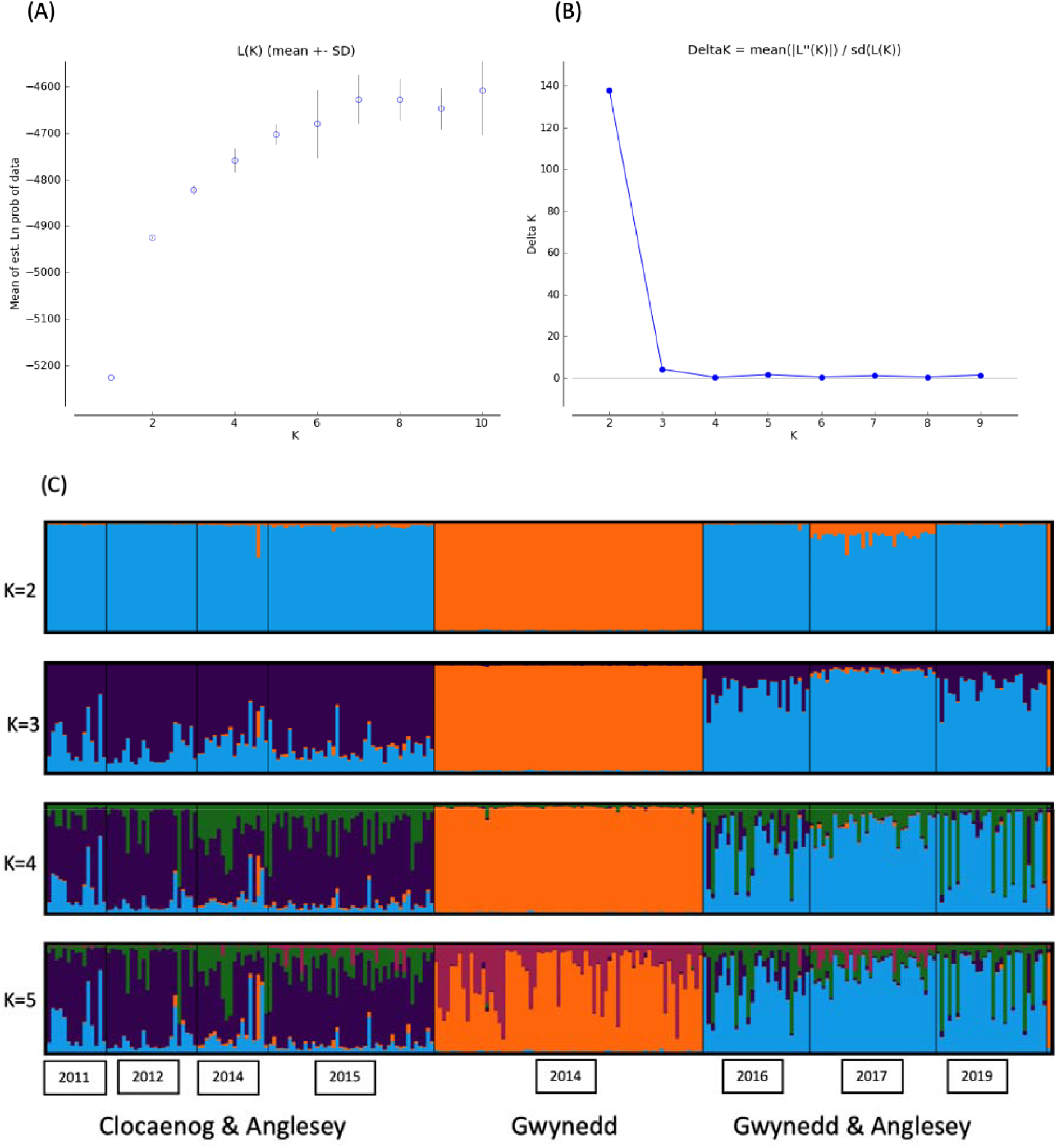
(A) plot of mean likelihood L (K) and standard deviation per *K* value of 254 individual grey squirrels genotyped at eight microsatellite loci. (B) Plot of Evanno’s ΔK method (Evanno et al 2005). (C) Membership of individual grey squirrels to K = 2, 3, 4 and 5 as inferred by STRUCTURE analysis. Each grey squirrel is represented by a single vertical bar. The geographic region and year of capture is indicated below the structure plot.

At K = 2, the majority of the animals clustered in one population, with a second cohort of animals culled in Gwynedd in 2014 forming the second cluster. Later culls in the Gwynedd area retained little evidence of this cluster, but an animal removed from Anglesey in 2020, also formed part of that 2014 cluster (Fig. 4C). Overall, there was evidence of genetic differentiation between Clocaenog and Gwynedd, with further substructure occurring within Gwynedd, particularly after the initial cull in 2014.

There was also some evidence of genetic clustering in the principal co-ordinates analysis (Fig. 5(A) and (B), which accounted for 20% of the genetic variation within the data.

**Figure 5.**
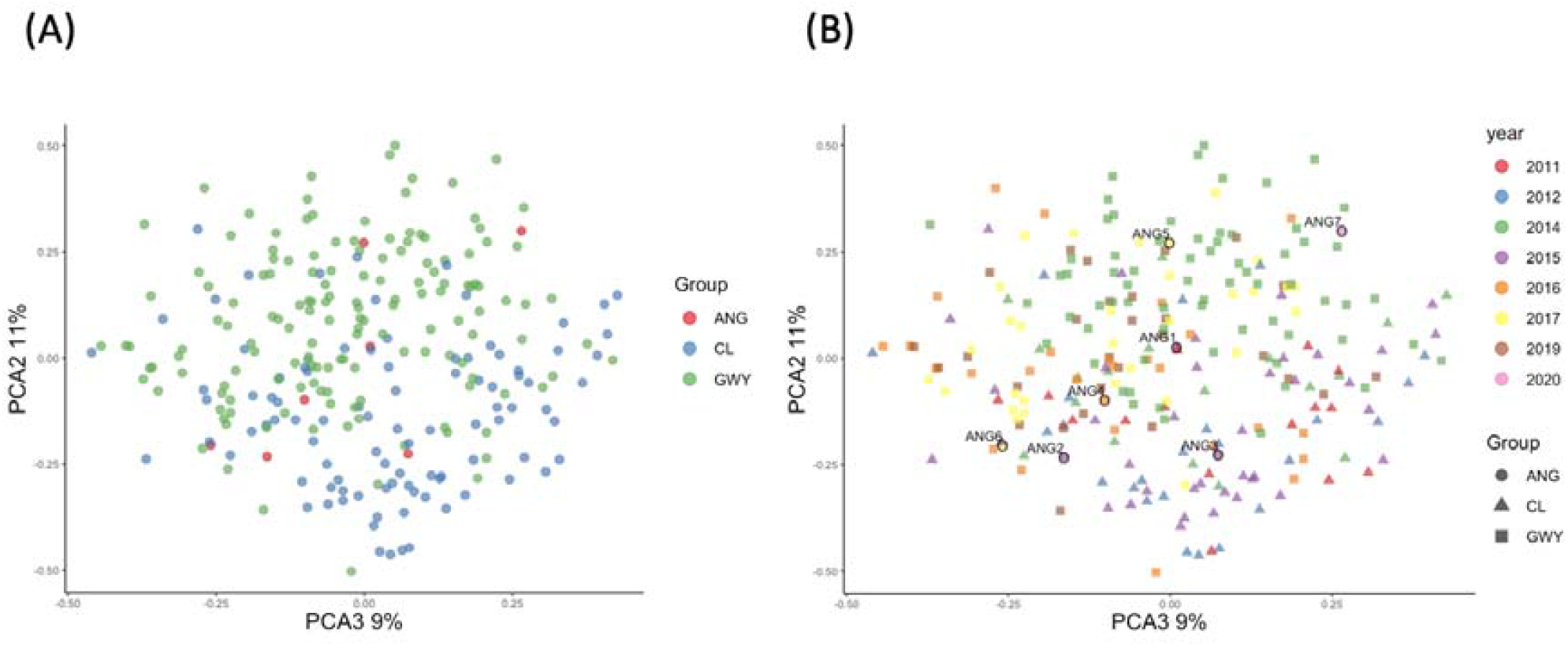
Principal coordinate analysis of individual red squirrels. (A) 20% of genetic variation of the data grouped by location, (B) 20% of genetic variation of the data grouped by location and year sampled (N = 254).

## Discussion

In this study, we demonstrated that genetic monitoring can be used as a method to evaluate the effectiveness of a long-term culling operation to reduce an invasive species population. We used a combination of mtDNA to gain historical insights into the introduction and establishment of the grey squirrel in north Wales, and temporal microsatellite DNA data to assess if culling had altered the genetic diversity. Previous studies have shown that culling is an effective management strategy as it shows a reduction in population size (Signorile & Shuttleworth 2016). However, our study suggests a scenario of relative population stability despite intensive control efforts. This is likely a consequence of the high number of historical introductions of genetically diverse grey squirrels from a broad geographical area in North America, and continued gene flow from neighbouring populations within north Wales containing genetically differentiated populations.

### mtDNA

A total of six mtDNA haplotypes were found within the north Wales population, three of which were new to this study. Twelve haplotypes were previously recorded throughout Britain, but these haplotypes were taken from a relatively small sample sizes; 14 from Stevenson et al (2013) and a further 73 from Stevenson-Holt & Sinclair (2015), and were representative of the populations in England and Scotland. However, given the high number of haplotypes found within north Wales alone, it is likely that the mtDNA diversity of the grey squirrel population is under recorded in Britain, and there are likely many other undocumented haplotypes present. Our mtDNA analysis showed that the haplotypes within north Wales are genetically diverse, and only two haplotypes, H9 and H15 positioned relatively close to each other on the network diagram (Fig. 3). Even the three haplotypes found on the island of Anglesey, H9, H12 and H14 are distinct from one another and likely originated from different introductions from North America. When comparing the British haplotypes to those recorded in North America (Moncrief et al 2012), the contemporary British population is highly diverse, with haplotypes from Britain found throughout the network which contained animals from its eastern and southern US distribution. Moncrief et al (2012) only sampled 69 animals from throughout this region, which again suggests that mtDNA within North America is also under recorded. However, despite the relatively low level of sampling, we found three haplotypes from North America in the contemporary British population, H8, H11 and H12. Haplotype H8 was previously recorded in Cumbria in northwest England, while H11 and H12 were recorded in Henbury in Northern England, a well-documented introduction site in Britain, from which other translocations took place (Stevenson-Holt & Sinclair 2015). The overlap of the H12 haplotype within all three studies can directly link grey squirrel movement patterns. This provides the information to trace the translocations from Maryland and Virginia in eastern North America to northern England and now to Wales. This possibly identifies a migration route of the grey squirrels from northern England to Clocaenog as this seems to be an entry point for moving populations into north Wales as it has the highest diversity of haplotypes in the areas sampled for this current study.

The high mtDNA diversity found within Britain today is testament to the large numbers of repeated introductions and translocations that took place over a 60-year period. This is further confounded by the species’ naturally high level of genetic diversity owing to its evolutionary history in North America where its genetic legacy appears not to have been imprinted by glacial history, limiting interpretation of its phylogeographic history (Moncrief et al 2012). Other *Sciurus* species have similar phylogeographic legacies including *S. niger* in North America and *S. vulgaris* in Eurasia (Grill et al 2009; Moncrief et al 2012; O’Meara et al 2018). All have a high number of mtDNA haplotypes that cannot generally be traced to commonly recognised glacial refugia. While this legacy may on the surface appear to be a contributing factor to the overall success of the grey squirrel in terms of its colonisation ability following repeated introductions, the same cannot be said for the red squirrel, who has also been reintroduced and translocated multiple times in parts of Britain and Ireland, and retains a similarly diverse genetic history (O’Meara et al 2018).

### Microsatellite

In relation to the microsatellite data, the grey squirrel population in north Wales is overall genetically diverse and contains relatively high levels of allelic richness (averaging 6.7) and high levels of expected heterozygosity (average H_E_ = 0.56), but rather low levels of observed heterozygosity (average H_O_ = 0.35) were obtained. The lower levels of observed heterozygosity are likely indicative of diminishing genetic diversity, which may also be related to the high levels of inbreeding evidenced within the population which were high and significant across all loci and averaged 0.35. The levels of allelic richness recorded in north Wales are higher than previously reported for Northumberland in northeast England (A_R_ = 4.3) and in East Anglia in the east of England (A_R_ = 3.44). Levels of expected and observed heterozygosity were also higher for both populations in Northumberland (H_E_ = 0.66; H_O_ = 0.64) and East Anglia (H_E_ = 0.71; H_O_ = 0.77), and less of a difference between both expected and observed values of heterozygosity were observed (Signorile et al 2014). F_IS_ values were also lower in both the Northumberland (0.04) and East Anglia (−0.15) populations (Signorile et al 2014) suggested that these populations are less inbred than populations in this current study. Given the high levels of inbreeding and the lower levels of observed heterozygosity in comparison to expected heterozygosity, it was anticipated that the population might currently or previously have experienced a genetic bottleneck. However, there was no evidence of a recent genetic bottleneck as evidenced by a normal ‘L’ shaped distribution of mode-shift test and non-significant heterozygote excess. It may be the case that the high levels of allelic richness are countering or protecting the population from entering a bottleneck. Bottlenecks are detected when levels of allelic richness decrease before a decrease in heterozygosity has occurred resulting in higher levels of heterozygosity than expected by the number of alleles (Luikart et al 1998). The high levels of inbreeding in absence of a genetic bottleneck is somewhat of a paradox, as invasive species are often expected to have overcome a genetic bottleneck during the invasion process (Frankham 2005), but in the case of the grey squirrel, it appears to have avoided a bottleneck, despite high levels of inbreeding, perhaps due to the high numbers of genetically diverse founders. Similar genetic legacies have been found in the invasive brown anole lizard (*Anolis sagrei*), which due to the effect of multiple introductions from different areas, resulted in higher genetic diversity in invasion sites than in its original native range (Kolbe et al 2004).

The impact of repeated culling over time appeared to illicit mixed results in relation to genetic diversity. For instance, observed and expected levels of heterozygosity generally decreased over time in the Gwynedd population between 2014 (H_O_ = 0.410; H_E_ = 0.612 and 2019 (H_O_ = 0.353; H_E_ 0.609), but actually increased in Clocaenog between 2011 (H_O_ = 0.336; H_E_ = 0.494) and 2015 (H_O_ = 0.372; H_E_ 0.551). However, variation also occurred within the sampling years, which is likely a consequence of geneflow and new animals entering the culled area, particularly in Clocaenog which is geographically closer to England and as was said earlier, is likely to have greater colonisation potential. This is further corroborated by an increase in the number of alleles and levels of allelic richness in the area, which actually increased during the culling period. Inbreeding levels were high and significant across all years sampled, and again no evidence of genetic bottlenecks were found. Indeed, the STRUCTURE and PCA results from this study have shown that the cull that took place in Gwynedd in 2014 contained a genetically distinct population, with further evidence of population differentiation occurring between later culls in Gwynedd and Cloceanog, suggesting a mechanism for how migrants can introduce new diversity into a population at arrival. It seems likely that the removal of animals creates a space whereby animals from nearby areas migrate to fill, and while Signorile et al (2016) found that grey squirrels exhibited relatively levels of migration, it seems that the removal or reduction of adjacent populations, encourages inward migration. In a conservation context, this mechanism is called a genetic rescue, whereby even a small number of new immigrants can contribute additional genetic variation to the remnant population reducing the risk of extinction (Whiteley et al 2014). In the case of the invasive grey squirrel, it may be a mechanism which supports and protects the genetic diversity of the species following intensive control efforts.

The animals that dispersed onto and were culled in Anglesey, although small in number, did not differ from the populations in Gwynedd or Clocaenog, and likely originated from both areas. The most recent animal culled in Anglesey in 2020 clustered with the animals culled from Gwynedd in 2014, suggesting that while this cluster appeared to have been removed by earlier culls, it is likely that it was not fully removed and may be recovering.

### Overall implications and recommendations

We processed grey squirrel samples that had been culled in north Wales between 2011 and 2020. This culling effort was costly both financially and in terms of manpower to attempt to remove grey squirrels from the woodlands of Gwynedd and Clocaenog and all surrounding areas, and our results have indicated that the population is not likely to be exterminated in the near future without sustained regular culling efforts. The likelihood of new animals moving into a culled area is also high, as we demonstrated in our microsatellite data analysis, that once an area was cleared of a population, a new and genetically distinct population moved in to occupy that empty space. The high levels of mtDNA diversity, even within a small area of north Wales adds further genetic variability to the population. The implications of this means that control efforts have to take place at regular intervals to ensure that new squirrels do not invade a cleared area, requiring further financial and labour supports.

Despite the broad ecological similarity and complete niche overlap between the invasive grey squirrel and the native red squirrels, it is notable that both having been introduced, reintroduced, and translocated multiple times in Britain, the red squirrel continues to be much more susceptible to the negative effects of a decrease in population size than the grey squirrel. However, the grey squirrel appears to be able to recover from a reduction in population size as it increases migration and gene flow. This ability appears to be one of the biological factors that makes the grey squirrel such a successful invasive species (e.g. Shuttleworth et al. 2020).

One of the ways invasive species are thought to be so successful is because they often do not encounter stress to the same degree that they do in their native environment and may be released from predatory pressures. This can result in an increase in fitness which can counter other eco-evolutionary processes normally experienced by species in their native environments such as inbreeding (Colautti et al 2017). Intriguingly, recent studies in Britain and Ireland have shown that the presence of a predator, the pine marten (*Martes martes*) (Sheehy & Lawton 2014, Twining et al 2020, 2021) is associated with a reduction of grey squirrel populations and thus facilitates a natural recovery of the red squirrel. As a result, a number of pine marten reinforcement projects have taken place in Britain (Sheehy et al 2018; McNicol et al 2020a, 2020b). Captive bred pine martens have been released in the Gwynedd area of our study (Bamber et al 2020). A combination of increased predator presence and an improvement in habitat may provide the best circumstances to maintain the Welsh red squirrel population, while actively reducing the grey squirrel. The investment in the restoration of predatory species like the pine marten to the north Wales countryside may provide benefits not only to the conservation and management of native and invasive species but will provide further increases in biodiversity benefiting the wider ecosystem and society in general.

## Acknowledgments

This project was financially supported by EU LIFE 14 NAT/UK/000467, The National Lottery Heritage Fund and Natural Resources Wales. The authors would like to thank Jack Bamber and Russell Layton for their contribution to the sample collection. We are also grateful to Dr Liz Halliwell and Iolo Lloyd for allowing access to historical samples collected in Clocaenog forest.

## Data Accessibility

The datasets generated during and/or analysed during the current study are available from the corresponding author on reasonable request. Raw sequence data will be made publicly available upon publication.

